# On the mechanisms of Transcranial Magnetic Stimulation (TMS): How brain state and baseline performance level determine behavioral effects of TMS

**DOI:** 10.1101/189969

**Authors:** Juha Silvanto, Silvia Bona, Marco Marelli, Zaira Cattaneo

## Abstract

The behavioral effects of Transcranial Magnetic Stimulation (TMS) can change qualitatively when stimulation is preceded by initial state manipulations such as priming or adaptation. In addition, baseline performance level of the participant has been shown to play a role in modulating the impact of TMS. Here we examined the link between these two. This was done using data from a previous study using a TMS-priming paradigm, in which, at group level, TMS selectively facilitated targets incongruent with the prime while having no statistically significant effects on other prime-target congruencies. Correlation and linear mixed-effects analyses indicated that, for all prime-target congruencies, a significant linear relationship between baseline performance and the magnitude of the induced TMS effect was present: low levels of baseline performance were associated with TMS-induced facilitations and high baseline performance with impairments. Thus as performance level increased, TMS effects turned from facilitation to impairment. The key finding was that priming shifted the transition from facilitatory to disruptive effects for targets incongruent with the prime, such that TMS-induced facilitations were obtained until a higher level of performance than for other prime-target congruencies. Given that brain state manipulations such as priming operate via modulations of neural excitability, this result is consistent with the view that neural excitability, coupled with nonlinear neural effects, underlie behavioral effects of TMS.

## Introduction

Single pulses of Transcranial Magnetic Stimulation (TMS) applied concurrently with a visual target can either facilitate or impair detection performance, depending on factors such as stimulation intensity and brain state. Whereas TMS intensities above phosphene threshold have been found to mask visual perception when applied over the early visual cortex within a time window of 80-120 ms from stimulus onset (see e.g. Kammer, 2005, de Graaf et al, 2014, for reviews), subthreshold TMS within the same time window can facilitate behavior (e.g. Abrahamayan et al, 2011), particularly when baseline performance is low, reflecting a situation with a weak perceptual signal (e.g. Schwarzkopf et al, 2011).

Nonlinear effects are also observed when TMS is applied during a behavioral task following an initial state manipulation such as adaptation or priming. For example, in a study using orientation-contingent colour priming, suprathreshold TMS (applied within the TMS-masking time window) was found to induce a *facilitatory* effect on items incongruent with the prime, with no effects on other prime-target congruencies (Silvanto et al, 2017). Similarly, TMS induces a facilitation of adapted visual attributes while the same stimulation parameters impair performance in the absence of adaptation (Silvanto et al, 2007). Thus manipulations of initial activation state qualitatively change the nature of behavioral TMS effects.

However, the extent to which these state-dependent TMS effects depend on the baseline performance level of the participant is unknown. This is a potentially important issue, given that, as discussed above, task difficulty has been shown to determine behavioral effect of TMS in conventional “virtual lesion” paradigms (e.g. Schwarzkopf et al, 2011). Furthermore, there is clear evidence that effects of another brain stimulation technique, transcranial direct current stimulation (tDCS), interact with baseline performance level (e.g. Tseng et al., 2012; Hsu et al., 2014; Juan et al., 2017). Thus one may query whether the facilitations observed in state-dependent paradigms be explained in terms of baseline performance level of the participants, such that TMS enhances performance of low performers but impairs high performers, as has been observed by prior TMS and tDCS studies (Schwarzkopf et al, 2011; Tseng et al., 2012; Hsu et al., 2014; Juan et al., 2017). We examined this issue by carrying out new analyses on our dataset from the state-dependent TMS study of Silvanto et al (2017), focusing on individual differences. In that study, participants were required to detect the colour of a briefly presented colour grating. On each trial, the target stimulus was preceded by a prime (a combination of colour and orientation) which was either fully congruent with the target (i.e. both colour and orientation matched), fully incongruent (ie. both colour and orientation of the prime and target differed), or partially congruent (i.e. either colour or orientation of the target matched that of the prime). Single pulse TMS was applied within the classic TMS-masking time window, 100 ms after target onset (e.g. de Graaf et al, 2014). The results showed that, at group level, single pulse TMS applied within the classic TMS-masking time window facilitated the detection of targets fully incongruent with the prime, while having no statistically significant effects on other stimulus types (see Figures 1). These effects were found on median reaction times of correct responses.

**Figure 1.**
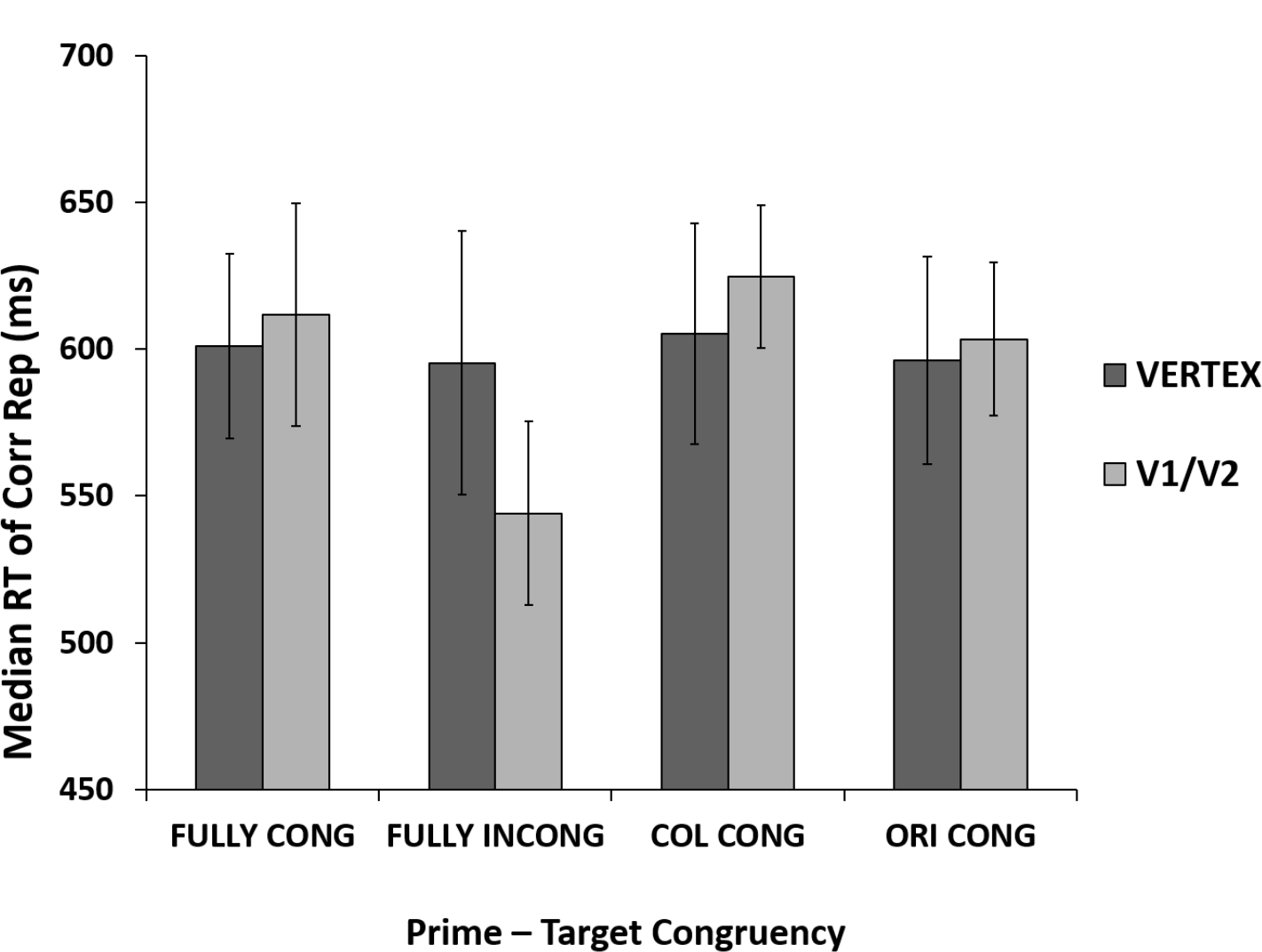
Results from Silvanto et al (2017). Statistical analyses showed a selective facilitation of fully incongruent trials. TMS had no significant effect on other prime-target congruencies.

To examine whether these effects are driven by or modulated by participants’ baseline level of performance, we carried out correlation and linear mixed-effects analyses to examine the relationship between baseline reaction times, priming manipulation and the induced TMS effect, as well as a new group analyses in which participants were divided into low and high baseline performance groups.

## Methods

The methods have been reported previously in Silvanto et al. (2017) and reproduced below for the reader’s convenience:

### Participants

33 participants (12 M, mean age=23.06 years) with normal or corrected-to-normal vision volunteered to participate in the experiment, of whom 18 perceived TMS-induced phosphenes and were thus included in the main analysis. One participant was excluded due to chance-level baseline performance. Thus analyses was carried out on 17 participants. All subjects provided written informed consent before participating in the study, which had been approved by the local ethics committee. All participants were naıve to the aims of the study and were treated according to the guidelines of the Declaration of Helsinki. Prior to participation, each participant was screened for contraindications to TMS.

### Stimuli and psychophysical task

Stimuli were presented at a viewing distance of 60 cm on a 16-inch monitor with a display resolution of 1920 × 1080. Stimuli and task were presented by using E-prime software (Psychology Software Tools Inc., Pittsburgh, PA). Both stimulus prime and stimulus target consisted of diagonal lines at 45° to the left or right of vertical such that stimuli were made of a series of stripes. These stripes were either black and green (CIE × = 0.30, y = 0.60, luminance 20 cd ⁄m2) or black and red (CIE x= ¼ 0.60, y = 0.35, luminance 20 cd⁄m2) with a stripe width of 0.25° in a stimulus subtending 6° horizontally and 3° vertically (adapted from Silvanto et al., 2007’s study). Therefore, four different colour–orientation combinations were used, in which prime and target could have: a) same colour and same orientation (“fully congruent” trial); b) opposite colour and opposite orientation (“fully incongruent” trial); c) same colour but opposite orientation (“colour congruent” trial); d) same orientation but opposite colour (“orientation congruent” trial). These congruency types appeared with equal frequency within a block. The procedure is shown in Figure 2. Each trial started with a fixation cross presented in the middle of the display for 500 ms, followed by the presentation of the prime stimulus (appearing for 100 ms) and subsequently by a 300ms blank screen; after that, the target stimulus appeared on the middle of the screen for 20 ms. The target stimulus was followed by a mask (remaining **on** the screen till participants’ response) composed of black diagonals in both possible orientations and with the gaps filled with green or red with the colour for each gap selected at random. A new randomly generated mask was used for each trial. When the mask was presented, participants had to indicate the colour of diagonals in the stimulus target display (red or green) by pressing the corresponding key on the keyboard. The prime lasted longer than the target in order to induce a stronger initial activation state manipulation. The logic was that a longer lasting prime will induce a stronger activation difference in primed vs non-primed neurons, and thus increase the likelihood of observing an interaction with TMS. The target needed to be of short duration so that the task is not trivial. Both accuracy and response speed were emphasized. Each participant underwent a total of 8 experimental blocks, namely 2 blocks for each stimulation site (V1/V2, Vertex) and for each stimulation intensity (80%, 120% of PT, see below). Each block included 40 trials, 10 for each of the 4 colour-orientation combinations. The order of stimulation sites and intensities was counterbalanced across participants, as well as the orientation–colour combinations of the stimuli within each block. Before the main experiment, participants underwent a block of practice with no TMS (20 trials).

**Figure 2.**
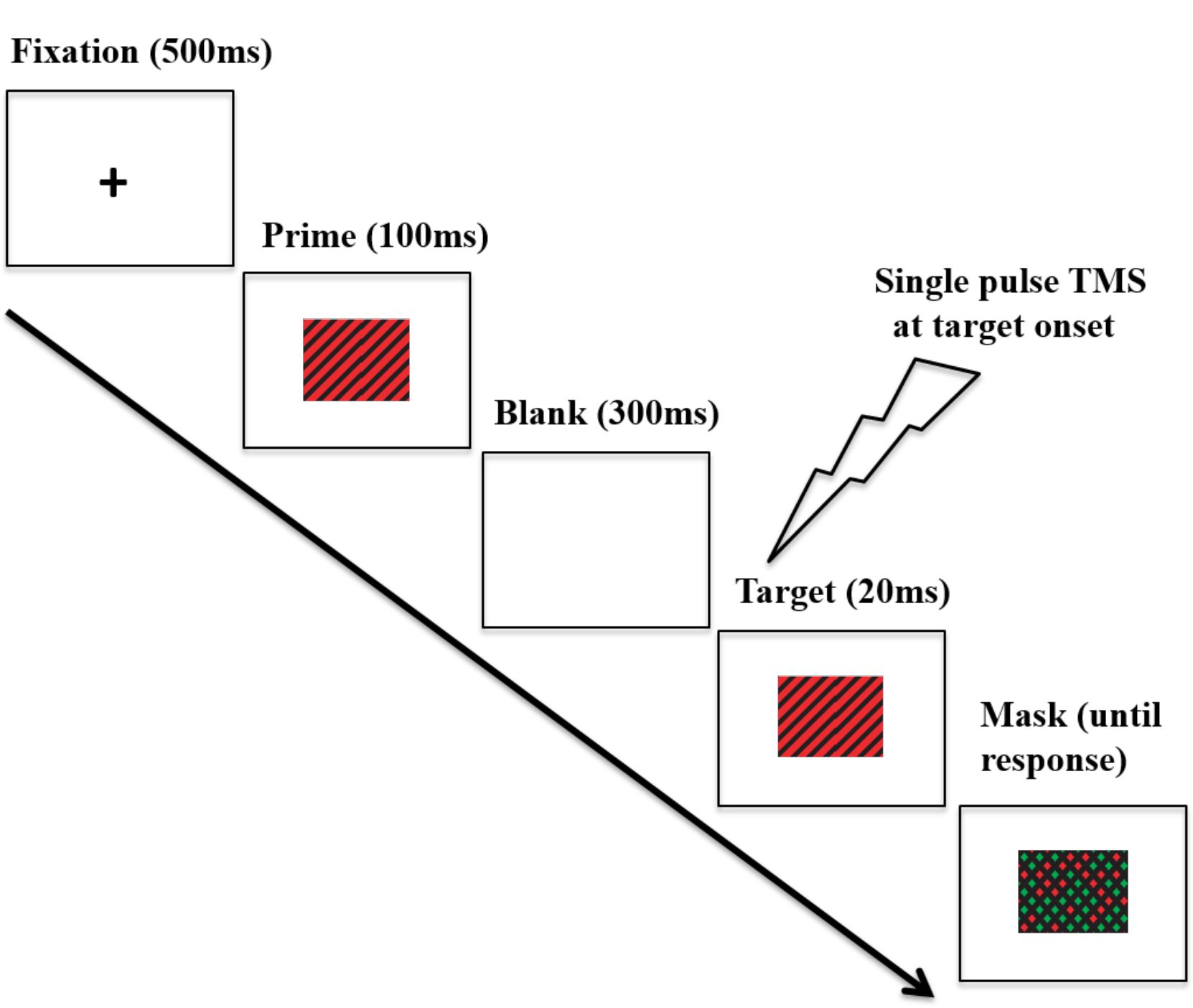
Timeline of an experimental trial. On each trial, participants were presented with a prime that was either a red-black or green-black grating, tilted either clockwise or counterclockwise. This was followed by a target which could be either fully congruent with the prime (i.e. the same stimulus), fully incongruent (i.e. both colour and orientation differed), or partially congruent (either colour or orientation matched the prime). Participants had to indicate the colour of diagonals of the stimulus target (red or green). In this figure, a fully congruent trial (i.e. prime and target matched for both colour and orientation) is depicted. Single pulse TMS was delivered at 100 ms after target onset over either V1/V2 region or over the Vertex (baseline). Adapted from Silvanto et al (2017).

### Transcranial Magnetic Stimulation

TMS was administered using a 70mm biphasic figure-of-eight coil connected to a Magstim stimulator (Magstim, Wales). The site of stimulation (V1/V2 region) was localized functionally in each participants by means of phosphenes search (see Walsh & Pascual-Leone, 2003, for a detailed description, see e.g. Campana et al, 2002, 2006; Cattaneo et al., 2011, for examples). In this method, the coil is initially positioned 2 cm above the inion and its location is subsequently adjusted until foveal phosphenes (overlapping with the target location in the main experiment) are induced. **Phosphene thresholds (PTs)** were measured, after dark adaptation, for each participant using a modified binary search algorithm (Tyrrell & Owens, 1988; Thilo et al., 2004). In the main experiment, participants were stimulated at 90% and 120% of their PT. On each trial, a single-pulse TMS was delivered over V1/V2 or Vertex (baseline), 100 ms after onset of the target stimulus, i.e. within the classic TMS masking time window (e.g. de Graaf et al, 2014). Vertex was identified as the halfway location between the inion and the nasion and at an equal distance from the left and right inter-trachial notches and was used as control site (Cattaneo et al, 2012; Bona et al, 2015). We included two Vertex conditions, one in which TMS was applied at 90% and the other with 120%, so that the level of possible auditory artefacts was controlled. During the stimulation, the coil was held with the handle pointing medial to lateral away from the midline and kept in place by the experimenter.

### Statistical Analyses

Statistical analyses were carried out on median reaction times of correct responses, as analyses on this variable showed statistically significant effects in Silvanto et al (2017), with a selective facilitation of fully incongruent targets while no effects were found on other stimulus types. Furthermore, we focused on the suprathreshold TMS intensity, as no TMS effects were observed with subthreshold intensity in Silvanto et al. (2017; see Figure 1).

Data were analysed as a function of congruency between the prime stimulus and the target – there were thus 4 trial types: congruent trials (i.e., prime and target are identical); incongruent trials (i.e., prime and target differ in both colour and orientation); colour congruent trials (i.e., target colour but not orientation matches the prime) and orientation congruent (i.e., orientation but not colour matches the prime).

## Results

### Correlation and linear mixed-effects analyses

We first examined the correlations between baseline (Vertex) level of performance (median RT of correct responses) and the magnitude of the induced TMS effect (defined as performance in baseline (Vertex) condition subtracted from the TMS condition). These are summarized in the scatterplots shown in Figure 3.

**Figure 3.**
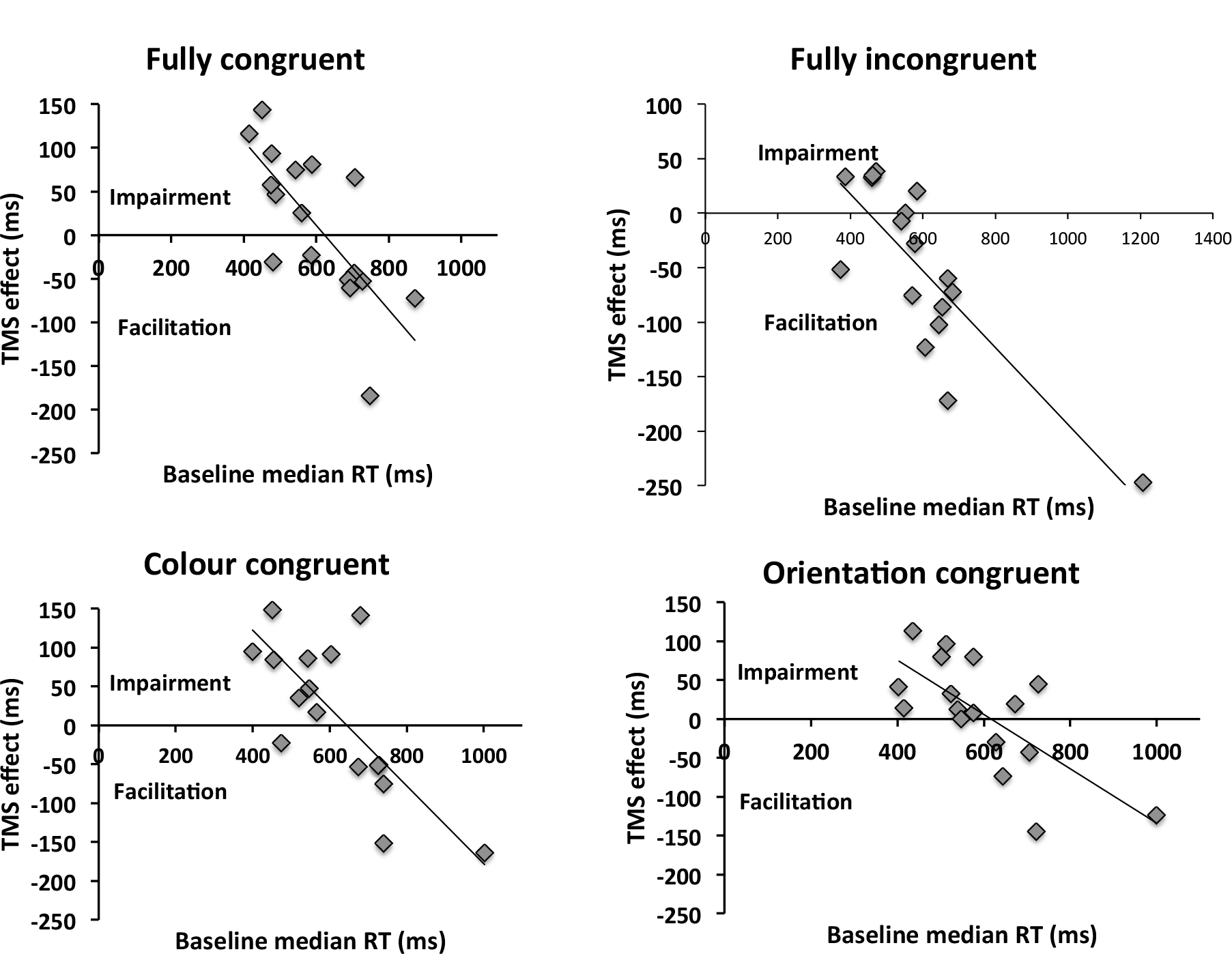
Relationship between baseline reaction time and the induced TMS effect. The TMS effect reflects the difference between V1/V2 TMS and Vertex TMS conditions. Values above the x-axis indicate impairment of behavior, whereas values underneath it reflect facilitation. A significant correlation was present for all congruencies. Incongruent condition appears to differ from other conditions in terms of the intersection with the x-axis, which reflects transition from inhibitory to facilitatory effects of TMS. This is in the region of 500 ms for congruent condition, whereas it is above 600 ms for other congruencies. The implication is that, for incongruent trials, facilitations are observed until higher levels of performance (i.e. at lower RTs) than for other congruency types. Thus, the facilitatory range of TMS effects is wider for the incongruent stimuli.

For all prime-target congruencies, correlation (Pearson’s r) between baseline performance and the TMS effect was significant (Fully congruent: r=−0.741, p=0.001; fully incongruent: r=−0.814; p<0.001; colour congruent: r=−0.772; p<0.001; orientation congruent: r=−0.703; p=0.002). These indicate a negative relationship between baseline level of performance and the TMS effect, such that low baseline performance is associated with a facilitatory effect of TMS and high baseline performance with impairments. A linear mixed-effects analysis was then performed to predict the TMS effect based on baseline RT and prime-target congruency (Baayen et al., 2008). By-participant random intercepts were included in order to account for the non-independency of observations. We employed the R packages lme4 (Bates et al., 2014) and lmerTest (Kuznetsova et al., 2015). To exclude the impact of overly influential outliers on the results, data points were removed on the basis of a threshold of 2.5 SD of the model standardized residual errors, and the model was then re-fitted (Baayen, 2008). The analysis showed significant impact on the TMS effect of both baseline performance (F(1,53.43)=95.95; p<.001) and prime-target congruency (F(3,40.57)=5.54; p=.003). However, no significant interaction between the two predictors was observed (F(3,40.65)=1.89; p=.146). The model-estimated regression lines are represented in Figure 4.

How does congruency modulate the TMS effect? The mixed-effects analysis reveals that the main difference between the different prime-target congruencies appears to be the intercept, which is affected by the main effect of condition. In fact, the intercept is significantly lower in the incongruent case (256 ms) relative to other congruency conditions (fully congruent 412 ms (t(80.63)=3.84; p<.001); colour congruent 426 ms (t(78.08)=4.79; p<.001); orientation congruent 347 ms (t(79.16)=2.49; p=.015)). In contrast, the slope (i.e., the effect of the baseline performance) is similar across conditions, as indicated by the non-significant interaction term (congruent: −0.669; incongruent: −0.518; colour congruent: −0.679; orientation congruent: − 0.571). The intercept parameters estimate the intersection points between each regression line and the y-axis (falling out of the plot areas in the Figures). However, as evident from Figure 3 and 4, because of the comparable slopes these different intercepts translate into different intersections with the x-axis, and hence with a difference in transition from facilitatory to inhibitory effects of TMS. This means that for the incongruent condition, the transition from TMS facilitating performance to impairing it, as a function of baseline performance level, occurs at a higher level of baseline performance (494ms, according to the prediction of the mixed effects model) vis-à-vis other conditions (fully congruent: 615 ms; colour congruent: 627 ms; orientation congruent: 607 ms). In other words, incongruent trials are facilitated until a higher level of performance than other trial types - effectively widening the facilitatory range of TMS effects.

**Figure 4.**
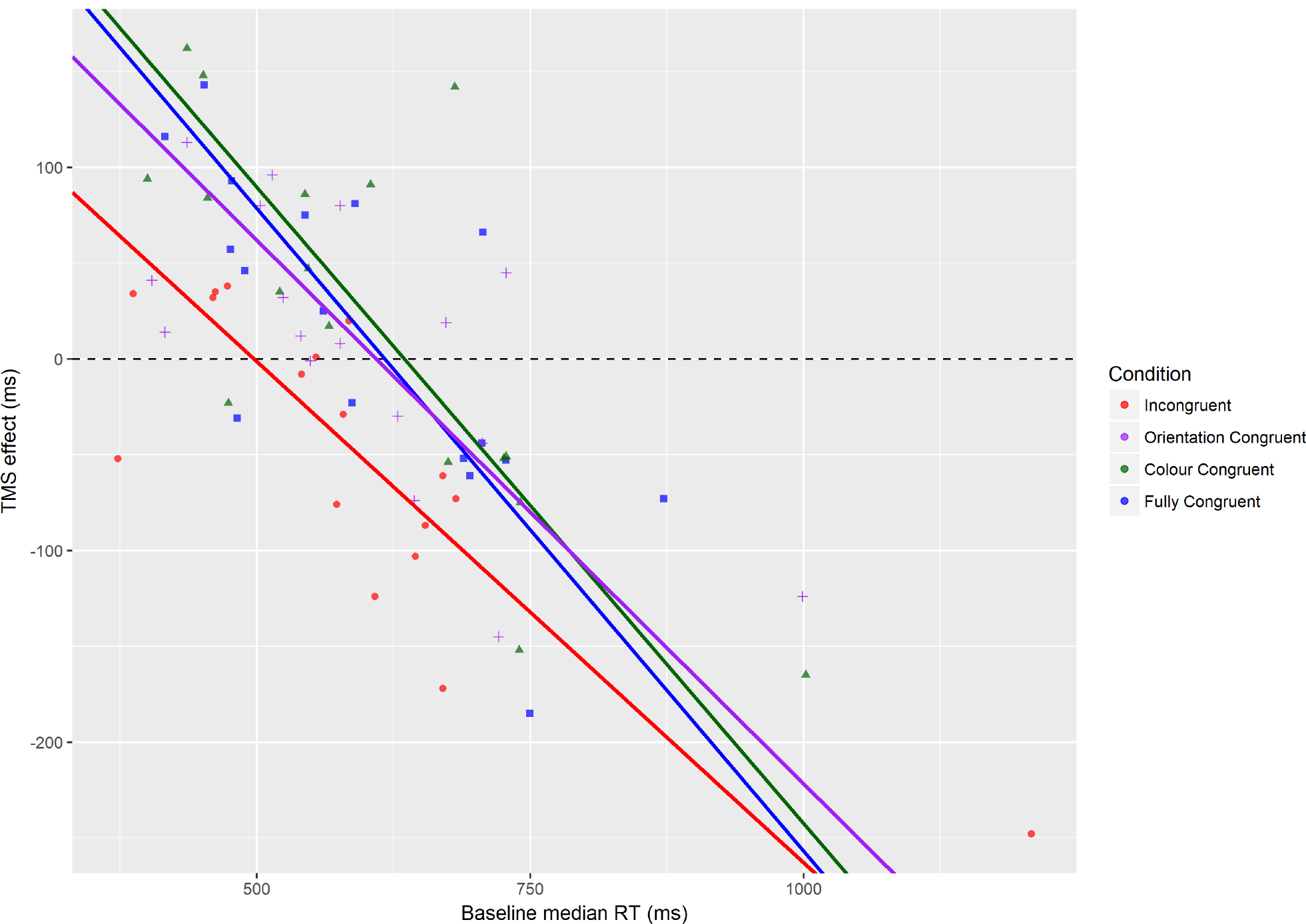
Regression lines for the different prime-target congruency conditions, as estimated by means of a linear mixed-effects model.

### ANOVAs on low and high baseline performers

The above analysis indicates that TMS may facilitate performance in the incongruent condition until a higher baseline performance levels than for other congruency types. To examine this further, we divided participants into two baseline performance groups (“low” and “high”). The results are shown in Figure 5 (see Figure 1 for the same results when data are not divided by baseline performance). For both groups, we carried out an ANOVA with congruency (fully congruent, fully incongruent, colour congruent, orientation congruent) and TMS site (Vertex, V1/V2) as within-subject factors.

**Figure 5.**
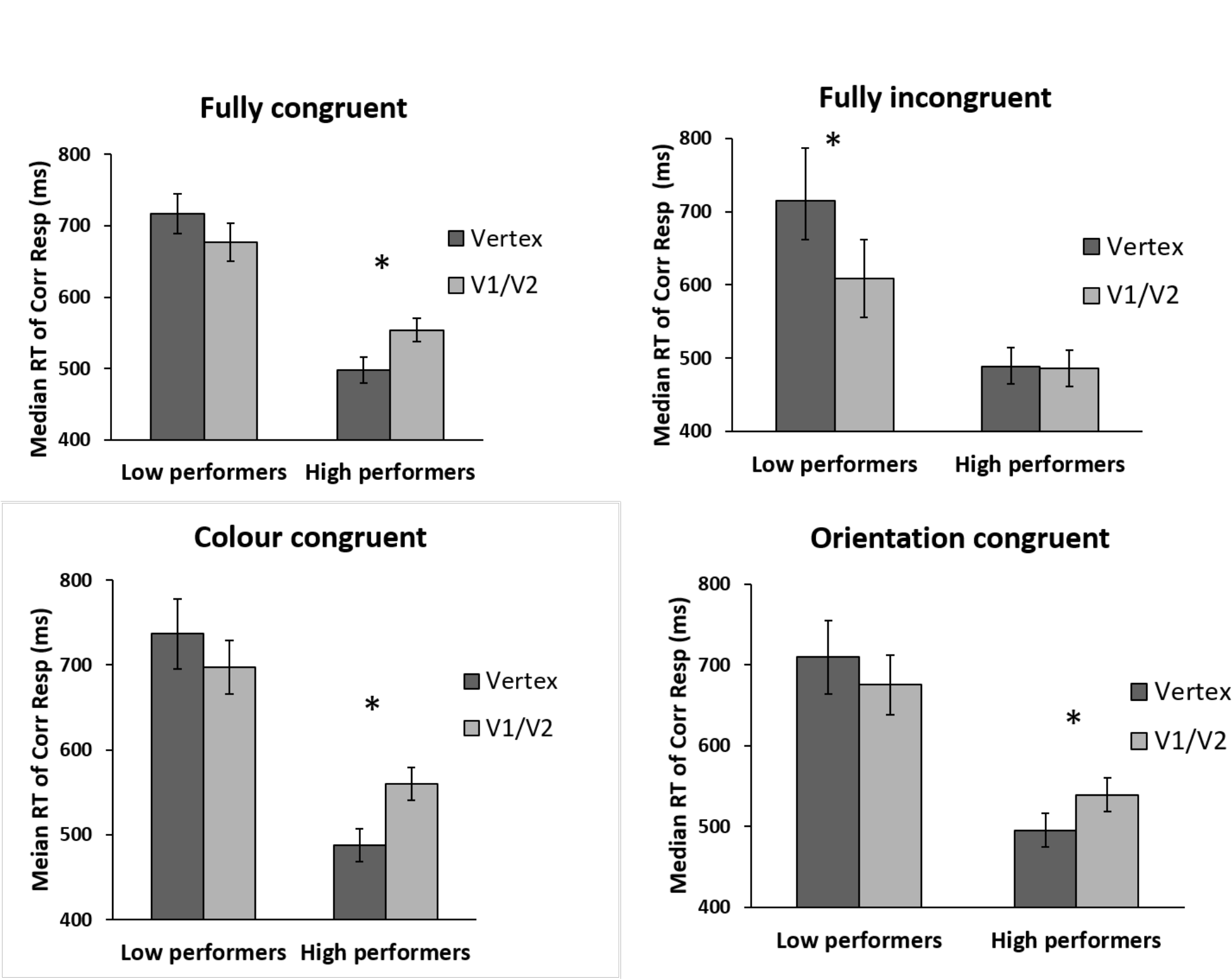
Performance of low and high baseline performers as a function of prime-target congruency and TMS site. For low performers, TMS induced a borderline-significant general facilitation regardless of prime-target congruency. For high performers, TMS impaired all congruency types expect incongruent trials.

For low *performers,* the main effect for TMS site was borderline significant (F(1,7)=5.107; p=0.058; η_*p*_^2^=0.422), with the main effect of congruency (F(3,21)=1.264;p=0.312; η_*p*_^2^=0.153) and interaction between site and congruency being non-significant (F(3,21)=2.293; p=0.11; η_*p*_^2^= 0.247). The borderline significant main effect of TMS indicates that the impact of V1/V2 TMS was to induce a weak general facilitation relative to Vertex TMS, regardless of congruency (see Figure 3). A planned t-test indicated that this facilitation reached significance only for the incongruent condition t(7)=3.766; p=0.007) – driving the effect reported in Silvanto et al. (2017). For other prime-target congruencies, this facilitation was not close to statistical significance (lowest p-value 0.21).

In contrast, *for high performers,* there was a significant main effect of TMS site (F(1, 8)=12.602; p=0.008; η_*p*_^2^=0.612), a borderline significant effect of congruency (F(3,24)=2.769; p=0.064; η_*p*_^2^=0.257) and a highly significant interaction between site and congruency (F(3,24)=5.068; p=0.007; η_*p*_^2^=0.388). Thus for high performers, congruency did significantly modulate the impact of TMS. Pairwise comparisons showed that V1/V2 TMS impaired performance relative to Vertex TMS for all other congruencies except for fully incongruent trials (*fully congruent:* t(8)=2.830; p=0.022); *colour congruent:* t(8)=3.611; p=0.007); *orientation congruent*: t(8)=3.120; p=0.014); *fully incongruent:* t(8)=−0.199; p=0.847).

In short, this analysis showed the following: for low performers, TMS induced a borderline-significant general facilitation regardless of prime-target congruency. (Planned pairwise comparisons indicated that for the incongruent trials there was a significant facilitation, driving the effect reported in Silvanto et al., 2017). For high performers, TMS impaired all congruency types expect incongruent trials.

## Discussion

Our findings indicate that baseline performance level and initial brain state combine to produce behavioral effects of TMS. Firstly, our results showed that a relationship between performance at baseline and the magnitude of the induced TMS effect was present for *all* prime-target congruencies. Thus, even though at group level TMS was found to modulate performance only for fully incongruent targets (see Silvanto et al, 2017), an analysis focusing on individual differences revealed the operation of a TMS effect at all levels of congruency. Specifically, there was a negative relationship between baseline performance level and the induced TMS effect, such that, as baseline performance level increased (i.e. RTs decreased), TMS effects turned from facilitatory to inhibitory.

Secondly (and what is the key finding), the *TMS effect depended on both condition and baseline performance.* In other words, both the baseline performance and prime condition contributed towards determining whether a facilitation or impairment was observed. Specifically, the incongruent condition differed from other congruency conditions in terms of the transition point from facilitatory to disruptive effects of TMS (reflected as intersection with the x-axis in Figure 3 and 4). As Figure 3 indicates, the effect of TMS was such that performance of participants with relatively slow baseline RTs was facilitated by TMS, whereas those with faster baseline RTs were impaired. Interestingly, for incongruent trials the transition from facilitatory TMS to inhibitory TMS was shifted relative to other congruency types. For incongruent trials, TMS-induced facilitation was present also for participants with a higher baseline performance, whereas facilitation in other conditions was only seen in participants with a lower baseline performance. Specifically, while for the incongruent condition the transition from facilitation to impairment occurred at approximately 500 ms, for other congruencies this occurred at lower performance levels, at approximately 600 ms. This shift in the intersection with the x-axis can explain why group-level analyses such as those carried out by Silvanto et al (2017) tend to reveal significant facilitatory effects only on incongruent trial types: as the “window” for facilitatory effects of TMS is wider (i.e. such effects are obtained with a large range of baseline performance), more participants will fall within this range. In contrast, for other congruencies the transition from facilitation to inhibition occurs with lower level of baseline performance, increasing the likelihood of observing group level *impairments* for these congruency types. The existence of a wider facilitatory vs. disruptive TMS window for incongruent vs other trial types is supported by the results of the ANOVA in which participants were divided into low and high baseline groups. In these analyses, TMS impaired high performers for all prime-target congruencies other than incongruent trials. In contrast, for low performers there was a borderline-significant main effect of stimulation site, indicating that TMS tended to facilitate performance for all stimulus congruencies; however, the facilitation was strongest for incongruent trials, due to the reasons discussed above.

Prior studies have shown that TMS can have a facilitatory effect on near-threshold stimuli (e.g. Abrahamyan et al, 2011, 2015; Schwarzkopf et al, 2011). The present results show that baseline performance level alone is insufficient to account for TMS effects, given that the same TMS parameters can have a different consequence despite similar baseline performance, as a function of brain state manipulations. While baseline performance (or “signal strength”) clearly plays a role, this does not provide the complete picture.

An important issue is that state manipulations such as priming modulate *neuronal excitability* (i.e. susceptibility to external input such as TMS). Behavioral priming is a phenomenon in which sensory systems are more efficient in processing stimuli which match those which have been presented previously. Effectively, priming modifies the amount of external stimulation required to activate neuronal populations (e.g., Kohn and Movshon, 2003; Kohn, 2007), such that an external stimulus of lower/higher strength is needed to drive the neurons tuned to primed/non-primed sensory stimuli (e.g. Gotts et al., 2012). Thus a lower TMS intensity is required to induce a similar neural effect in primed neurons, relative to non-primed ones (see Silvanto et al, 2017, for further discussion).

Therefore, priming is likely to reduce excitability of neural representations incongruent with the prime and therefore a higher TMS intensity is needed to drive these neurons. How can this explain the shift in facilitatory/inhibitory range found here? We have previously proposed (see Silvanto et al, 2017; Silvanto & Cattaneo, 2017) that behavioral TMS effects can be explained in terms of changes in neural excitability interacting with nonlinear neural effects, as a function of TMS intensity. The latter have been reported by Moliazde et al. (2003), who found that low intensity TMS induced a facilitation in neural activity and visually-induced neural firing lasting up to 200 milliseconds, followed by longer lasting neural suppression. In contrast, with high TMS intensities the early facilitation was replaced by a suppression of neural activity. This early facilitation vs. inhibition of neural activity has been linked to behavioral facilitations vs. impairments, which also occur at low vs. high TMS intensities (i.e. at group level, subthreshold TMS facilitates and suprathreshold TMS impairs perception (see Silvanto et al, 2017; Silvanto & Cattaneo, 2017, for more detailed discussions of this view). The key point is that, changes in neural excitability, by modifying susceptibility to TMS, shift the intensities with which facilitatory and inhibitory effects of TMS are observed. This occurs because the same TMS parameters have a stronger or weaker effect when excitability has been increased/decreased. In a simplified sense, reduction in excitability (as is the case here with incongruent trials), turns inhibitory *high* intensity TMS to facilitatory *low* intensity stimulation - as the same stimulation intensity has a weaker neural effect after priming. The consequence is a shift in the transition point from facilitatory to disruptive effects of TMS for incongruent trials.

In this view, why does baseline performance level matter? It matters because baseline reaction times can be seen as a measure of neural excitability, in terms of the efficacy with which the incoming stimulus is processed by the visual system. Slower reaction times are indicative of lower excitability, reflecting slower information accumulation before perceptual threshold is reached. According to the model of Silvanto & Cattaneo (2017), lowering of neural excitability increases the likelihood of a facilitatory TMS effect being observed, when TMS is applied at suprathreshold intensities. This is the case because, as discussed above, reducing excitability is akin to reducing TMS intensity, which turns “inhibitory” high intensity TMS to “facilitatory” low intensity stimulation. In contrast, fast reaction times are indicative of high excitability, and therefore more likely to fall within “inhibitory” range of TMS effects.

What are the implications of these findings for TMS studies more generally? Fundamentally, our results reflect the importance of conceptualising TMS effects as an interaction between neural excitability and TMS intensity, with the outcome depending on the interaction between the two. A given TMS intensity may either impair or facilitate behaviour, depending on the strength of the external stimulus and the readiness of the perceptual system to encode that stimulus. It is important to note that such nonlinearities are observed not only in state-dependent TMS paradigms, but are likely to be general feature of brain stimulation studies (e.g., Schwarzkopf et al, 2011; Tseng et al., 2012; Hsu et al., 2014; Juan et al., 2017). For cognitive neuroscientists using brain stimulation to study brain-behaviour relations, the implication is that observing a TMS effect (or lack of an effect) at any given intensity or perceptual/cognitive state may not be the whole story; in standard experiments, we generally observe only a specific combination of excitability and stimulation intensity – and the observed effect may very well be different at other combinations. In addition, it is also important to acknowledge that the timing of TMS in priming paradigms modulates the induced effect (Chiau, 2017). Thus to fully exploit the potential of TMS to characterise how a given brain area processes perceptual information, one should assess performance at different levels of baseline performance, timing and TMS intensity.

